# Dominance hierarchies are linear but shallow in the social amoeba *Dictyostelium discoideum*

**DOI:** 10.1101/2024.07.15.602882

**Authors:** Cathleen M.E. Broersma, Elizabeth A. Ostrowski

## Abstract

Social groups often form dominance hierarchies, and these hierarchies are almost always linear. However, why linear dominance hierarchies emerge is not well understood. In the social amoeba *Dictyostelium discoideum*, cells form a multicellular fruiting body when starved, which consists of a ball of viable spores held aloft by a stalk of dead cells. In genetically mixed (‘chimeric’) fruiting bodies, conflicts can arise over the equitable sacrifice of cells to the dead stalk, and some strains predictably dominate others in the spores. Using pairwise mixes of strains that co-occurred in small soil samples, we determined the dominance hierarchies in four natural populations of *Dictyostelium*. These hierarchies were significantly linear in two of four populations, but also extremely shallow, indicating that co-occurring strains are competitively similar. We used quantitative genetic analyses to assess the causes of social dominance. Each strain’s solo spore production was a significant predictor of its performance in pairs. However, we detected additional genetic contributions of both the focal and partner strain, indicating additional cryptic traits that mediate social competitiveness. In contrast to earlier studies showing strong fitness differences among strains collected over a larger spatial scale, we show that co-occurring strains are remarkably competitively equivalent, resulting in linear yet shallow hierarchies. Our results underscore the importance of biologically relevant spatial scales in assessing fitness interactions among microbes. They also explain why social trait diversity might be observed despite dominance hierarchies that should eliminate this variation.

## Introduction

Social interactions within a group frequently result in the formation of a dominance hierarchy, commonly referred to as a “pecking order” (Schjelderup-Ebbe 1922). These social structures emerge from repeated antagonistic interactions between individuals that result in the establishment of a clear dominant and subordinate individual (Drews 1993). For example, fish and rodents establish dominance through aggressive behaviors that involve chasing and biting, whereas crustaceans exhibit claw-grasping (Chase et al. 2002). Based on the outcomes of the individual dominance relationships, a ranking of all individuals in the group is established. An individual’s rank can significantly impact their fitness, with higher-ranked individuals obtaining greater access to limited resources and reproductive opportunities compared to lower-ranked individuals (Wilson 1975; Clutton-Brock, Albon, and Guinness 1984; Nelissen 1992; Goessmann, Hemelrijk, and Huber 2000).

The structure of a dominance hierarchy can be characterized by two properties: its linearity and steepness (de Vries, Stevens, and Vervaecke 2006). These properties are assessed by analyzing the collective of pairwise, or ‘dyadic’, interactions within the group (Drews 1993; de Vries 1995; de Vries, Stevens, and Vervaecke 2006). In a perfectly linear hierarchy, one individual dominates all others, the second dominates all but the most dominant, and so on. For example, in a triad of individuals A, B, and C, the hierarchy is linear if the relationships are transitive, e.g., if A>B, B>C, and A>C, where “>” indicates dominance (Figure 1A). In contrast, a hierarchy is increasingly non-linear when one or more intransitive triads exist, e.g., if A>B, B>C, but C>A. The steepness of a dominance hierarchy in turn reflects the differences in dominance ability between adjacently ranked individuals, with a hierarchy being shallow when the differences are small and steep when the differences are large (Figure 1B) (Vehrencamp 1983; Barrett et al. 1999; de Vries, Stevens, and Vervaecke 2006).

**Figure 1.**
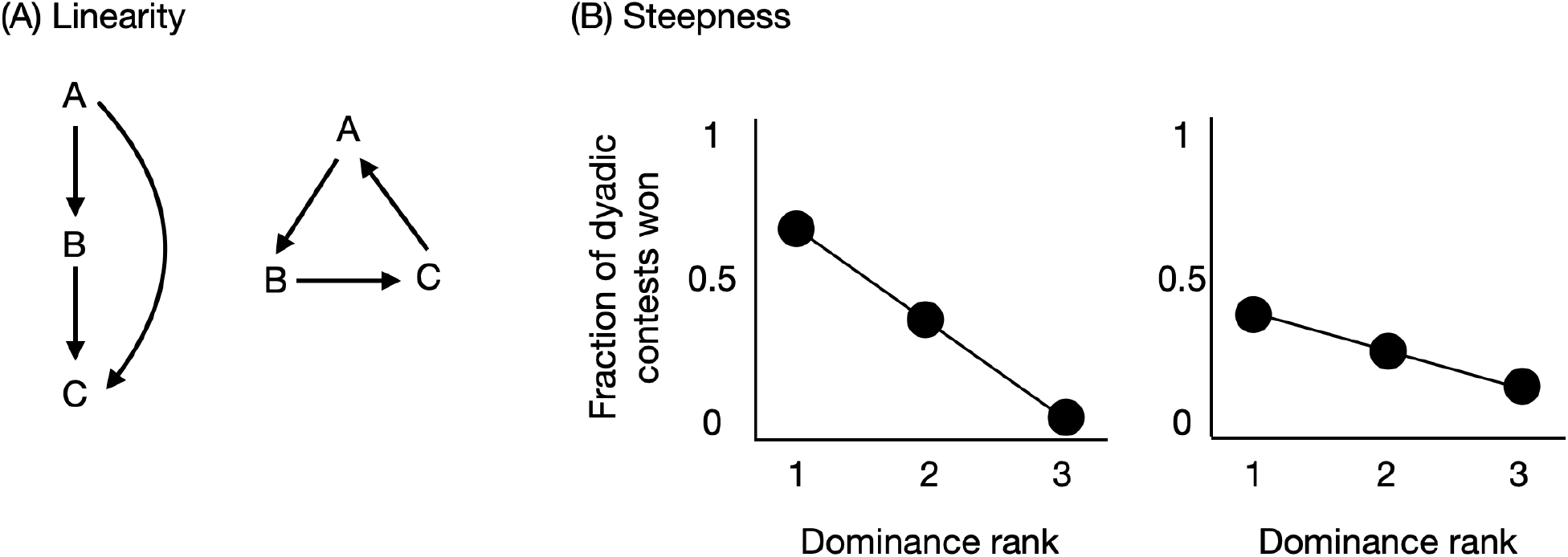
Two properties of a dominance hierarchy. (A) Linearity. In a perfectly linear dominance hierarchy, all triads are transitive (e.g., A>B, B>C, and A>C; left side). Alternatively, in a less linear dominance hierarchy, one or more triads are nontransitive (e.g., A>B, B>C, but C>A; right side). (B) Steepness. A dominance hierarchy is steep if the differences between adjacently ranked individuals are large and outcomes are highly repeatable (left). A hierarchy is flat if the differences between adjacently ranked individuals are small and/or less repeatable (right).

The majority of dominance hierarchies examined so far have been linear (Hausfater, Altmann, and Altmann 1982; Barkan et al. 1986; Heinze 1990; LeBrun 2005; Valderrabano-Ibarra, Brumon, and Drummond 2007; Wittemyer and Getz 2007; Correa et al. 2013; Vullioud et al. 2019). One major hypothesis to explain this linearity proposes that differences in individuals’ pre-existing attributes predict their success in social contests (Chase et al. 2002; Chase and Seitz 2011). Consequently, the hierarchy can be predicted before it has formed, and this scenario is therefore referred to as the ‘prior attributes’ hypothesis (Jackson and Winnegrad 1988; Drews 1993; Chase et al. 2002). These prior attributes may relate to physical (size and weight), personality (aggressiveness), genetic (genotype), social (maternal rank), and physiological (hormone levels) characteristics (Jackson and Winnegrad 1988; Drews 1993; Nakano and Furukawa-Tanaka 1994; Beaugrand and Cotnoir 1996; Goessmann, Hemelrijk, and Huber 2000; Sundström et al. 2004; Schjolden, Stoskhus, and Winberg 2005; Chase and Seitz 2011). However, identifying which dominance attributes are of significant influence on dominance and quantifying their relative importance for the establishment of a hierarchy and its linearity remains challenging (Appleby 1983; de Vries 1995; Chase and Seitz 2011; Shizuka and McDonald 2012; Schmid and de Vries 2013).

The social amoeba *Dictyostelium discoideum* serves as an effective model system for investigating social dominance and dominance hierarchies. In response to starvation, thousands of unicellular amoebae aggregate to form a multicellular fruiting body, which consists of a stalk of nonviable cells that lifts and supports a sorus of viable spores (Huss 1989; Kessin 2001). The self-sacrifice of the stalk cells is considered to be an altruistic act, as it likely benefits the survival and dispersal of the spores (Kuzdzal-Fick et al. 2007; Smith, Queller, and Strassmann 2014). Importantly, when genetically different individuals aggregate to form a single “chimeric” fruiting body, some strains dominate by preferentially forming spores and relying on the stalk formation by their social partner, such that spore production is unfair (Filosa 1962; Hamilton 1964a, 1964b; Buss 1982; Strassmann, Zhu, and Queller 2000). Consequently, chimeric fruiting body formation can involve a dominance relationship with a dominant and subordinate strain, offering a basis for constructing a dominance hierarchy among a group of co-occurring strains (Fortunato, Queller, and Strassmann 2003).

In two prior studies that examined the dominance hierarchy within a group of natural strains of *D. discoideum*, both revealed linearity (Fortunato, Queller, and Strassmann 2003; Buttery et al. 2009). In support of the prior-attributes hypothesis, Buttery *et al*. (2009) showed that pre-existing differences in spore production between strains during clonal development could predict social dominance and linearity in the group. However, both studies used the same seven strains, thus establishing a linear dominance hierarchy only once. They also found a slightly different order of strains, suggesting that the dominance hierarchies may not be repeatable. Moreover, these strains were collected within a 1 km^2^ area at a single site in North Carolina. Owing to the large spatial scale over which these strains were sampled, it is unknown whether the strains interact in nature and, therefore, whether the observed dominance hierarchy is ecologically relevant. Indeed, strains collected over more relevant spatial scales might be expected to exhibit lower dominance levels and less linear hierarchies. In other words, co-occurring strains might co-occur *because* they exhibit non-transitive relationships. Thus, assessing the linearity of dominance hierarchies over more realistic spatial scales is a priority.

Here we examined the dominance relationships among strains in four natural populations of *D. discoideum*. The strains that make up a population were isolated from soil taken from the surface of a 10-by-10 cm plot. Thus, the maximum distance between strains at the time of collection was 14 cm (i.e., the diagonal of the square plot).

Interactions between strains at this distance are reasonable, given that *D. discoideum* slugs routinely travel multiple centimeters from their aggregation sites (e.g., see Jack et al. (2015)), and moderate gene flow occurs over distances <0.5 km (Kuzdzal-Fick, unpublished). From the observed dominance relationships, we constructed and evaluated the linearity and steepness of the dominance hierarchy. For those populations that did show significant linearity, we asked if pre-existing differences in dominance ability, assessed through the clonal spore production, could predict the dominance outcome and a strain’s rank in the hierarchy. Additionally, we performed a quantitative genetic analysis to estimate the genetic basis of variation in social dominance. This analysis examined a focal strain’s success as a function of its own genes (‘direct genetic effects’, or DGEs), those of its partner (‘indirect genetic effects’, or IGEs), and their interaction (<u>g</u>enotype <u>x g</u>enotype (G×G) epistasis). IGEs and G×G epistasis are collectively referred to as the social environment, and existing studies have demonstrated that these factors are widespread and can significantly alter evolutionary trait dynamics compared to expectations based solely on DGEs (Moore, Brodie, and Wolf 1997; Wolf et al. 1998; Bijma and Wade 2008).

Of the four natural populations of *D. discoideum* we tested, two exhibited a significantly linear dominance hierarchy. Conversely, the other two populations did not show significant linearity, with one being marginally nonlinear. Notably, in the cases where linearity was observed, the dominance hierarchies were shallow, indicating minimal fitness differences among co-existing strains, at least in terms of their ability to dominate in the spores in a chimera. The lack of linearity in two populations, combined with shallow dominance hierarchies in all populations indicates co-occurring strains that are well-matched for social fitness. Lastly, the quantitative genetic analyses showed the importance of genetic identity and traits of the focal individual but also that of its social partner on the level of social dominance of the focal individual. These results demonstrate the importance of the social genetic environment and demonstrate that linearity can emerge even when social dynamics are present.

## Materials and Methods

### Strain collection and selection

The strains of *D. discoideum* used in this study were acquired between 2016 and 2019 by staff and students of the Ostrowski laboratory. Soil sampling and strain isolation methods are described in detail by Kuzdzal-Fick *et al*. (2023). The strains and sampling sites are listed in Table S1 and shown in Figure 2A.

**Figure 2.**
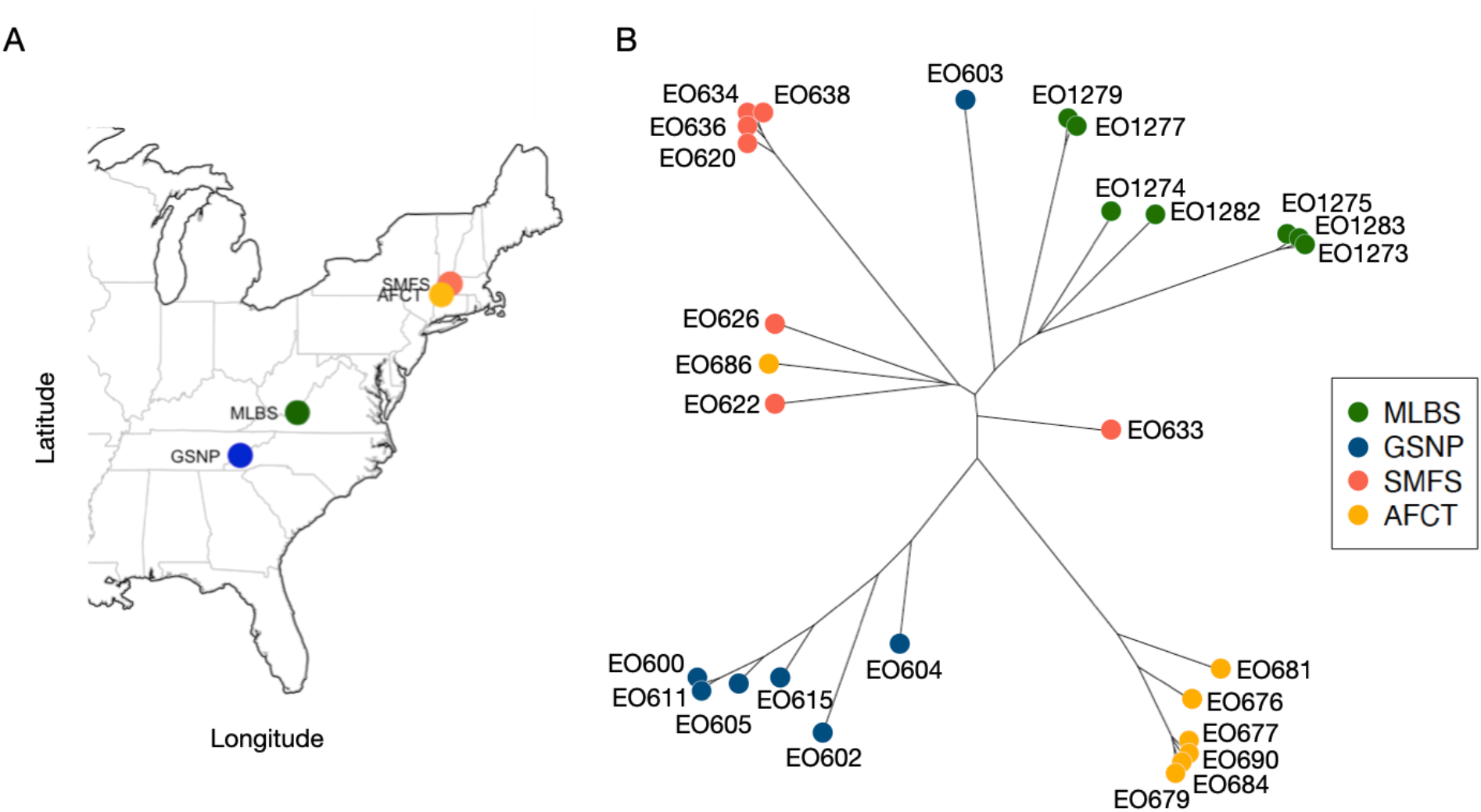
**(A) The 27 strains of *D. discoideum* in this study were sampled from four sites across the US. (B) A distance-based tree showing the genetic similarity among the strains based on genome-wide SNPs.**

To ensure that the chosen strains were not clones of one another, we used Illumina genome sequencing to determine the pairwise genetic distances among the strains from each site. Illumina sequence data was provided by Kuzdzal-Fick, and detailed information and methods concerning this data are described in Kuzdzal-Fick *et al*. (2023). We chose strains with a Euclidean genetic distance of >5 (or 1,756 SNPs), as our analyses indicated that values below this cut-off are within the margin of error for clones for this sequencing methodology. Figure 2B shows a genetic distance-based tree based on the Euclidean distance.

### Experimental design

We examined all possible pairwise combinations of strains for each of the four sites. For three sites with seven strains each —MLBS, GSNP, and AFCT— this resulted in 21 mixes. For site SMFS, with six strains, this resulted in 15 mixes. The lower number of strains in site SMFS was a result of having excluded strain EO631 because it did not progress beyond the slug stage during the experiments. We tested all combinations of strains from GSNP and SMFS on the same day, and from MLBS and AFCT on a different day. We performed three temporally independent blocks, each initiated from frozen spore stocks.

### Cultivation, cell staining, and developmental assays

At the start of each block, we inoculated the relevant strains from the freezer onto SM-agar plates (Formedium Ltd, 2% agar) with *Klebsiella pneumoniae* as a food source. After fruiting bodies had formed, we collected and replated 5×10^5^ spores with *K. pneumoniae* on fresh SM-agar plates for germination and population expansion. After approximately 40 hours, we harvested the cells during the mid-exponential growth phase and washed them three times via differential centrifugation (450 x *g* for 3 min) in KK2 buffer (14.0 mM K_2_HPO_4_ and 3.4 mM KH_2_PO_4_, pH 6.4) to remove the bacteria. After washing, the cells were resuspended to a density of 1×10^7^ cells/ml in KK2.

To distinguish the two strains in a mix, we treated one strain in each pair with the fluorescent dye CellHunt Green CMFDA (Molecular Probes), diluted in DMSO according to the manufacturer’s specifications. We added the cell dye at a concentration of 20 µM, incubated the cells on a shaking platform in the dark for 30 minutes, washed them once in KK2, and incubated them again for 30 minutes to allow the efflux of excess dye. We then washed the cells twice in cold KK2 and resuspended them at a concentration of 1×10^8^ cells/mL in cold KK2. We treated the unlabeled strain with DMSO only, and these samples underwent the same treatment as the labeled strains.

For each mix, we combined equal volumes of the labeled and unlabeled cells and deposited a 50 µl aliquot of the mix in a 3-by-3 square (an area of 1 cm^2^) of a 47 mm gridded 0.45 µm nitrocellulose filter, resulting in a total cell number of 5×10^6^ cells (Figure 3). We placed the nitrocellulose filter in a 6-cm Petri dish on top of a Pall filter moistened with 1.5 ml of PDF (per litre: 1.5 g KCl, 1.07 g MgCl·6H_2_O, 1.8 g KH_2_PO_4_, 1.6 g K_2_HPO_4_, 0.5 g streptomycin sulphate). We transferred the Petri dishes to a sealed plastic box that contained wet tissues and placed the box in the dark for 48 hours at 22°C to allow fruiting bodies to develop.

**Figure 3.**
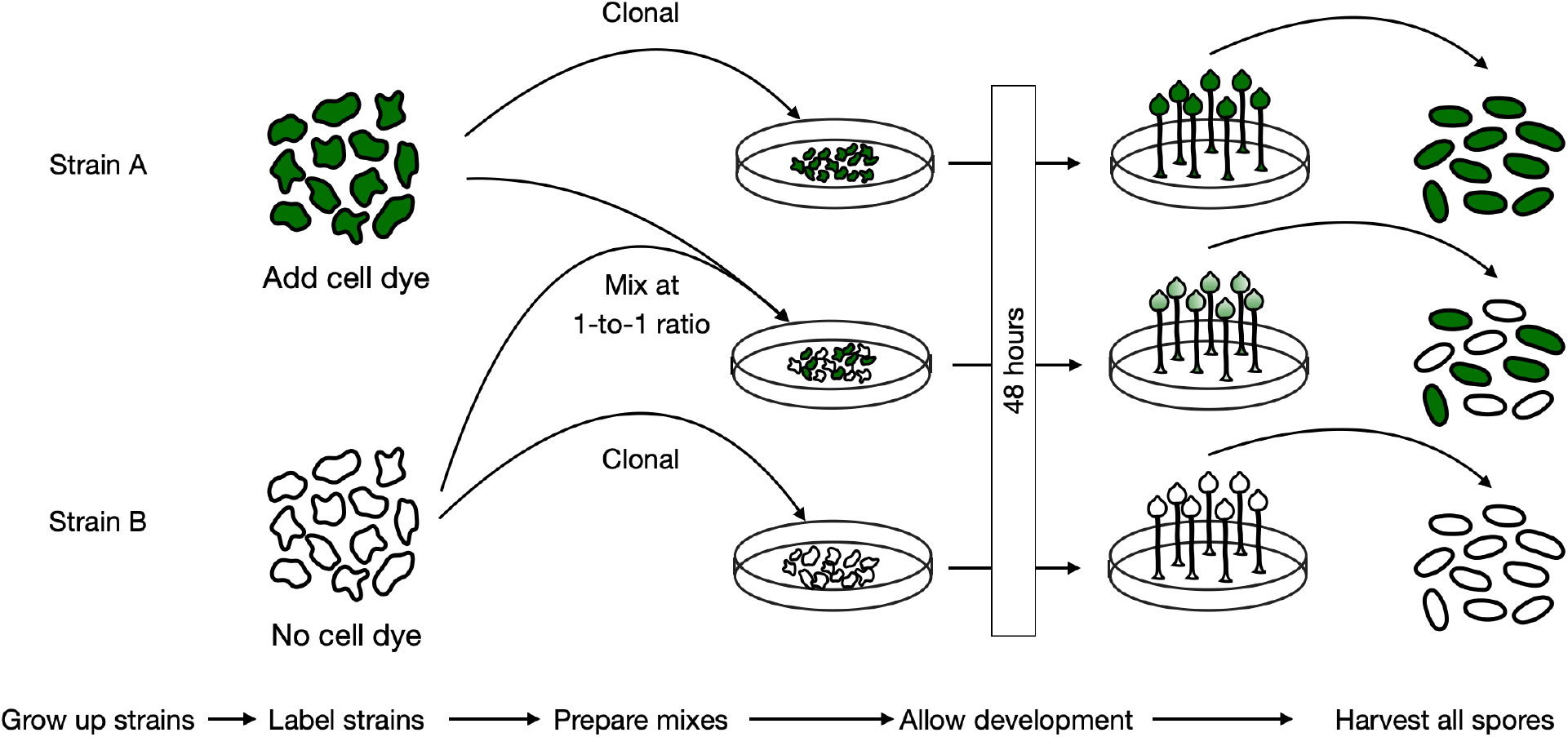
Overview of the mix experiments to assess social dominance. For each mix, the cells of a labeled and unlabeled strain were combined at equal cell numbers, spotted on a nitrocellulose filter, and allowed to develop fruiting bodies. After development, the spores were harvested, and the fractions of the labeled and unlabeled strains were determined using a flow cytometer. For each mix, the labeled and unlabeled cells of both strains were developed by themselves, as well as in a ‘control mix’ consisting of labeled and unlabeled cells of the same strain.

Following the development, we collected the spores in 3 mL of detergent (KK2 buffer + 0.1% IGEPAL and 10 mM EDTA), which dissolved any remaining cells. We measured the fraction of both strains in the spores on a BD FACSCanto II flow cytometer (488 nm laser, 513/15 GFP filter). We used an automated cell counter (Cell Countess II FL, Thermo Fisher) to quantify the total number of spores. For each mix, we developed the labeled and unlabeled cells of each strain by themselves. We also developed mixes that consisted of labeled and unlabeled cells of the same strain, which we refer to as the ‘control mix’ (Figure 3). All three types of clonal controls (100% labeled, 100% unlabeled and the 50-50% control mix) were otherwise treated identically to the chimeric mixes.

### Determination of the winner and loser strain and quantification of social dominance

For each mix, we determined the ‘winner’ and ‘loser’ strain based on their average fraction in the spores (across *N*=3 blocks). Since the strains were combined at equal starting cell ratios, we identified the winner as the strain with >50% of the spores, and the loser as the strain with <50% of the spores. We quantified the winner’s extent of social dominance as the magnitude of spore inequity: the increase in the strain’s representation in the spores compared to its initial representation in the cells. This measure ranges from 0 to 0.5, where a value of near 0 indicates roughly equal fractions of the winner and loser in the spores, and a value of 0.5 indicates that the winner makes up all the spores. For example, a social dominance value of 0.1 indicates that the winner makes up 60% of the spores in the chimera (i.e., increased by 10% relative to its starting frequency).

### Quantification of the linearity of the dominance hierarchy

We calculated the linearity of the dominance hierarchy using standard methods (Appleby 1983). Briefly, we constructed a win/loss matrix of *N*-by-*N* individuals, with the order of the individuals initially being arbitrary. We assigned an entry in the matrix a value of 0 if the row individual was the ‘loser’ when mixed with the column individual (<50% of the spores), and a value of 1 if the row individual was the ‘winner’ when mixed with the column individual (>50% of the spores). Using the row sums (*S*_*i*_) derived from the matrix, we calculated the observed number of transitive triads (*d*) using the formula:

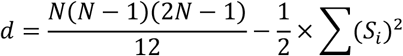

We calculated the degree of linearity (*K*) using the formulas:

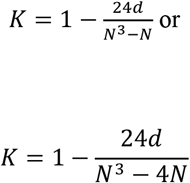

for even or uneven values of *N*, respectively. To assess the significance of *K*, we determined the degrees of freedom (*df*) and chi-square (*χ*^2^), since the distribution of *d* approaches that of a chi-square distribution with increasing *N*. The formulas for these calculations are:

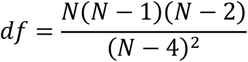

and

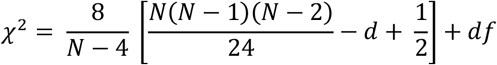

In the given formula, when the group size is *N*=6, the df equals 30, and when *N*=7, the df equals 23.3. According to a chi-square reference table, *K* was considered significant at *P*<0.05 if *χ*^2^>43.8 when *df*=30 and if *χ*^2^>35.2 when *df*=23.3.

In response to observing a very low estimate of steepness (discussed in the Results), we also calculated the triangle transitivity using the ‘transitivity’ function from the ‘EloRating’ package. Briefly, this function calculates the proportion of transitive triangles (*P*_*t*_) using the formula:

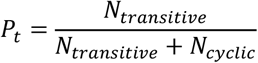

in which *N*_*transitve*_ is the number of transitive triads in the group, and *N*_*cycline*_ is the number of intransitive triads (Shizuka and McDonald 2012). The triangle transitivity (*t*_*tri*_) was then calculated as:

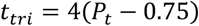

*t*_*tri*_ is scaled to run from 0 to 1, where 0 indicates the random expectation (0.75; regardless of the *N*) and 1 indicates maximum transitivity. Statistical significance of *t*_*tri*_ was assessed via randomization. A *P*<0.025 or *P*>0.975 means that the hierarchy is significantly more or less linear than expected by chance, respectively, based on 1,000 randomizations that were used to determine the null-distribution.

### Quantification of the hierarchy steepness

The hierarchy steepness quantifies the absolute differences in dominance ability between consecutively ranked individuals in a hierarchy. Steepness therefore also serves as a measure of ranking certainty. To calculate hierarchy steepness, we used the ‘steepness’ function from the ‘EloRating’ package (Neumann et al. 2011). In contrast to the binary win/loss matrix that we used to calculate the linearity, steepness was calculated using continuous data—specifically, the fraction of a strain in the spores. Steepness values range from 0 to 1, where 0 indicates a shallow hierarchy with small differences between adjacent ranks, and 1 indicates a steep hierarchy with large differences between adjacent ranks, and higher-ranked individuals consistently win against adjacent lower-ranked individuals (Figure 1B). The null hypothesis is that the observed steepness is no greater than a steepness based on random win probabilities (de Vries, Stevens, and Vervaecke 2006). To test this, randomization is used. Here the statistical significance (right-tailed *P*-value) of the observed steepness is determined by calculating the proportion of times that the steepness generated randomly under the null hypothesis is greater than or equal to the observed steepness (based on *N*=1000 iterations).

### Determining the ranking of strains in a hierarchy

For those populations that demonstrated significant linearity, we obtained the strain ranking using the ‘isi13’ function from the ‘compete’ package (Curley 2016). This function uses randomization to identify the optimal order of strains in a linear hierarchy (based on *N*=1000 iterations). The resulting linear order is that with the fewest inconsistencies (*I*) and the smallest inconsistencies (*SI*) among all potential orders. An inconsistency describes the situation where a lower-ranked individual dominates a higher-ranked individual in the assumed linear order (Schmid and de Vries 2013).

### Do strains vary in their prior attribute clonal spore production?

To assess if strains within a site vary in their inherent ability to dominate, we used a generalized linear mixed model (GLMM) that examined the clonal spore production as a function of strain ID and block. Because the response variable was a count, we used a Poisson error structure with a log link function. We included both terms as random effects and assessed their significance through single deletions of terms, comparing the reduced models with the full model using a likelihood ratio test.

### Does linearity arise from differences between strains in their ability to produce spores?

To test the hypothesis that linearity arises from a pre-existing hierarchy based on differences between individuals in their inherent ability to sporulate, we first ranked strains within each site based on their clonal spore production. The strain that produced the most spores clonally was ranked first, and so on. We then compared this ‘prior attribute ranking’ with the ranking based on the observed dominance relationships determined from the mixes above. The expectation, under the prior attribute hypothesis, was that the rankings in the two hierarchies would align.

### Quantitative genetic effects analysis

We performed two GLMMs to investigate the effects of clonal spore production, the genotypes of the focal and partner strain, and the interaction between them, on the focal strain’s relative (fraction; Model 1) and absolute (total; Model 2) number of spores in a mix.

In Model 1, we used beta regression since the response variable was a fraction between 0 and 1. Given the paired data (with fractions summing to 1), we randomly selected one strain from each pair as the focal strain and ran the model on this subset. We modelled the *clonal spore production* of the focal and partner strains and assessed their significance through type II Wald chi-square tests. We modelled the *genotype* of the focal strain, the partner strain, and their interaction as random effects. In addition, we included block as a random effect. We assessed the significance of the random effects through single deletions of terms, comparing the reduced models and the full model using a likelihood ratio test. Finding a significant effect of the focal genotype would indicate the presence of direct genetic effects (DGEs), a significant effect of partner genotype would indicate the presence of indirect genetic effects (IGEs), and a significant effect of the interaction between the focal and partner genotype would indicate the presence of genotype-by-genotype (G×G) interactions, or epistasis. To quantify the proportional contributions of the DGE, IGE and G×G interaction terms, we divided the variance components of each random term by the sum of the variance components of all random terms.

Model 2 incorporated several modifications compared to Model 1. Specifically, as the response variable in Model 2 was a count (i.e., the number of spores of the focal strain) and showed overdispersion, we used a negative binomial error structure (family=nbinom2). In addition, we included genetic distance between two strains in a chimera as a covariate and we used the full data set because the total spore production of both strains in a pair was independent. We included mix_id as a random variable to correct for including the spore production of both strains in a mix. We analyzed all other terms and performed tests of significance as described in Model 1.

In summary, with these models, we aimed to *i*. assess if the prior attribute ‘spore production during clonal development’ serves as a significant predictor of a strain’s relative (Model 1) and absolute (Model 2) dominance over spore production during chimeric development, and *ii*. quantify the relative importance of direct genetic effects (DGEs), indirect genetic effects (IGEs) and genotype-by-genotype (G×G) interactions on the focal strain’s spore production during chimeric development.

## Results

### Dominance hierarchies are mostly linear but shallow

Tables 1-4 show the dominance matrices for the sites AFCT, GSNP, MLBS, and SMFS, respectively. Each matrix value indicates the average spore fraction obtained by the strain in that row when co-developed with the strain in the corresponding column (based on *N*=3 blocks). Table 5 compares the linearity and steepness estimates for the four sites, along with the previously studied North Carolina (NC) site by Fortunato *et al*. (2003). Fortunato’s study, which tested all pairwise combinations of seven strains collected from Little Butt’s Gap (NC), demonstrated significant linearity, though steepness was not determined. We calculated the steepness for the NC hierarchy using their reported values (Table 2 from Fortunato *et al*. 2003). Similar to the NC site, sites AFCT and MLBS showed significant linearity based on Kendall’s *K* metric (Table 5). Sites GSNP and SMFS did not show significant linearity. Steepness for each hierarchy, including NC, did not significantly differ from zero. To address potential issues that may result from calculating linearity at a low steepness, we additionally calculated the triangle transitivity (*t*_*tri*_) (Sánchez-Tójar, Schroeder, and Farine 2018). Using this method, we observed a lower degree of linearity, but the hierarchies of sites AFCT and MLBS remained significantly linear (Table 5). In addition, using this metric, the hierarchy in site GSNP was marginally nonlinear (observed value is in the 3.4^th^ percentile of the null distribution—i.e., *P*=0.068; explained in the Methods).

**Table 1-4.** The dominance hierarchy matrix for sites SMFS, AFCT, MLBS, and GSNP respectively. Each entry indicates the mean fraction of the spores achieved by the strain in that row when co-developed with the strain listed in the corresponding column, based on *N*=3 blocks.

| *SMFS* | EO620 | EO622 | EO626 | EO633 | EO634 | EO638 |
| --- | --- | --- | --- | --- | --- | --- |
| EO620 | - | 0.526 | 0.515 | 0.489 | 0.488 | 0.518 |
| EO622 |  | - | 0.417 | 0.513 | 0.461 | 0.443 |
| EO626 |  |  | - | 0.388 | 0.527 | 0.481 |
| EO633 |  |  |  | - | 0.514 | 0.520 |
| EO634 |  |  |  |  | - | 0.477 |
| EO638 |  |  |  |  |  | - |

| *AFCT* | EO676 | EO677 | EO679 | EO681 | EO684 | EO686 | EO690 |
| --- | --- | --- | --- | --- | --- | --- | --- |
| EO676 | - | 0.631 | 0.596 | 0.586 | 0.475 | 0.450 | 0.424 |
| EO677 |  | - | 0.479 | 0.505 | 0.478 | 0.391 | 0.335 |
| EO679 |  |  | - | 0.532 | 0.544 | 0.508 | 0.409 |
| EO681 |  |  |  | - | 0.498 | 0.458 | 0.461 |
| EO684 |  |  |  |  | - | 0.446 | 0.493 |
| EO686 |  |  |  |  |  | - | 0.505 |
| EO690 |  |  |  |  |  |  | - |

| *MLBS* | EO1273 | EO1274 | EO1275 | EO1277 | EO1279 | EO1282 | EO1283 |
| --- | --- | --- | --- | --- | --- | --- | --- |
| EO1273 | - | 0.450 | 0.493 | 0.478 | 0.389 | 0.470 | 0.461 |
| EO1274 |  | - | 0.527 | 0.533 | 0.413 | 0.537 | 0.512 |
| EO1275 |  |  | - | 0.485 | 0.405 | 0.469 | 0.442 |
| EO1277 |  |  |  | - | 0.417 | 0.394 | 0.464 |
| EO1279 |  |  |  |  | - | 0.475 | 0.610 |
| EO1282 |  |  |  |  |  | - | 0.509 |
| EO1283 |  |  |  |  |  |  | - |

| *GSNP* | EO600 | EO602 | EO603 | EO604 | EO605 | EO611 | EO615 |
| --- | --- | --- | --- | --- | --- | --- | --- |
| EO600 | - | 0.577 | 0.537 | 0.628 | 0.495 | 0.476 | 0.501 |
| EO602 |  | - | 0.504 | 0.623 | 0.572 | 0.421 | 0.478 |
| EO603 |  |  | - | 0.623 | 0.572 | 0.421 | 0.478 |
| EO604 |  |  |  | - | 0.502 | 0.506 | 0.467 |
| EO605 |  |  |  |  | - | 0.515 | 0.485 |
| EO611 |  |  |  |  |  | - | 0.603 |
| EO615 |  |  |  |  |  |  | - |

**Table 5.** Summary of the properties of the four dominance hierarchies from this study and the NC population tested in Fortunato *et al*. (2003). *K* is a metric of linearity, ***P***_***t***_ is the proportion of transitive triads, and ***t***_***tri***_ is the triangle transitivity. See main text for explanations of terms. For site GSNP, *t*^*tri*^ is strongly negative and in the 3^rd^ percentile of the distribution based on randomization (i.e., 966/1000 randomizations were more positive than the observed value, and thus *P*=0.034*2=0.068 (the *P*-value is doubled because significance is now tested in the opposite direction, i.e., two-tailed test).

| | $K$ | $\chi^2$ | df | $P(K)$ | $P_t$ (expect 0.75 if random) | $t_{tri}$ | $P(t_{tri})$ | steepness | $p_{steepness}$ |
| --- | --- | --- | --- | --- | --- | --- | --- | --- | --- |
| SMFS | 0.250 | 38.7 | 30 | <0.90 | 0.700 | -0.200 | 0.777 | 0.030 | 1.00 |
| AFCT | 0.786 | 40.0 | 23.3 | <0.025 | 0.914 | 0.657 | 0.029 | 0.064 | 1.00 |
| MLBS | 0.929 | 45.3 | 23.3 | <0.005 | 0.971 | 0.886 | 0.009 | 0.060 | 1.00 |
| GSNP | 0.143 | 16.0 | 23.3 | <0.90 | 0.657 | -0.371 | 0.966* | 0.040 | 1.00 |
| NC (Fortunato *et al.* 2003) | 0.929 | 45.3 | 23.3 | <0.005 | 0.971 | 0.886 | 0.009 | 0.304 | 0.36 |

### Are dominance hierarchies generally linear in populations of D. discoideum?

The North Carolina (NC) site, as tested by Fortunato, exhibited the same linearity as site MLBS (Table 5; *K*=0.929, *χ*^2^_23.3_=45.3, *P*<0.005). Despite this similarity, MLBS displayed a very shallow hierarchy, with a steepness approaching zero, a pattern also observed in AFCT, GSNP, and SMFS. Notably, all four steepness values were substantially lower than that of the NC population. The lack of steepness in our study indicates much lower social dominance and, therefore, more equitable spore production. Specifically, in the Fortunato study, the average social dominance was 0.26 (across *N*=21 mixes), indicating a shift from a 50-50% distribution in the cells to 76-34% in the spores. In contrast, in this study, the average social dominance was only 0.045 (across *N*=78 mixes; Figure 4). Significant social dominance was observed in only 9% of all mixes: in 3 out of 21 mixes from GSNP, 2 out of 21 mixes from AFCT and MLBS, and none of the 15 mixes from SMFS. Significance was assessed based on a one-sample t-test, where the null hypothesis is that the observed social dominance is equal to that of the control mixes (Figure 4). In those mixes where we observed significant social dominance, the magnitude was 0.094±0.011 (mean±se, *N*=7), indicating a shift from a 50-50% distribution in the cells to 59-41% in the spores.

**Figure 4.**
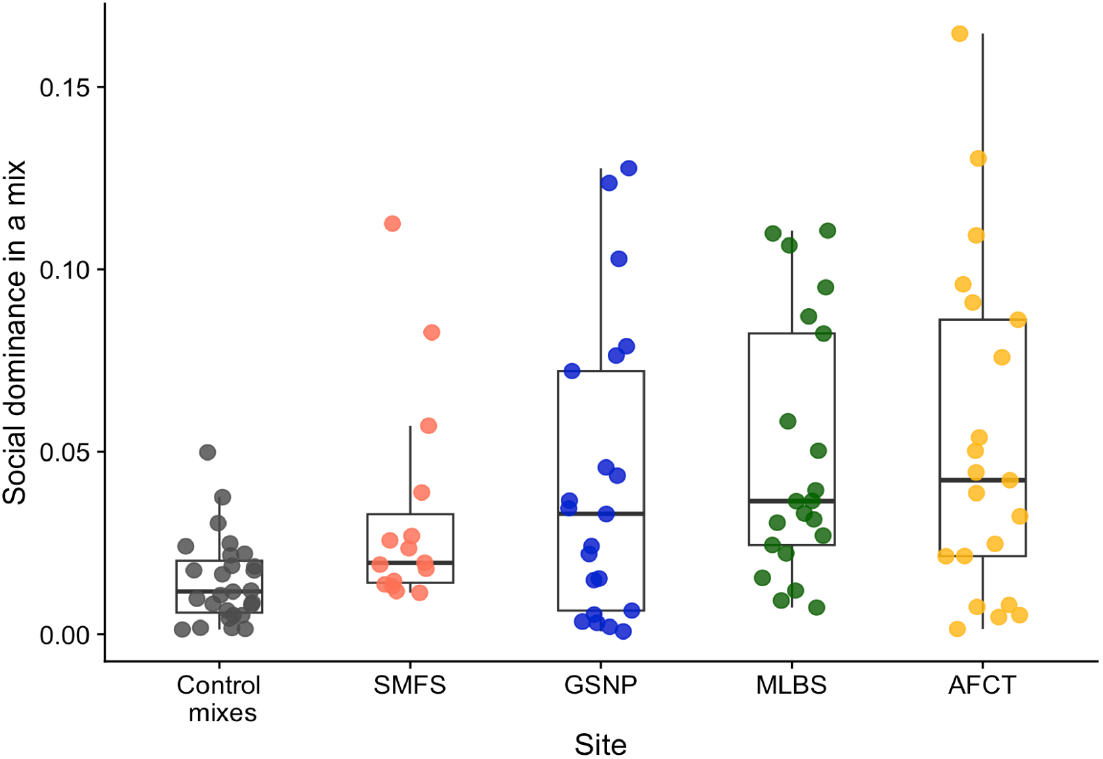
The magnitude of social dominance in control mixes and mixes of genetically different strains. Each point indicates the mean social dominance in a mix across *N*=3 blocks. For example, a value of 0.10 indicates a shift of 10%: from 50-50% in the cells to 60-40% in the spores. The control mixes are ‘mixes’ that consist of labeled and unlabeled cells of the same strain; these mixes serve as a baseline against which the mixes of genetically different strains can be compared.

### Can differences in the inherent ability to dominate in the spores explain linearity?

Prior work found that social dominance and linearity resulted from inherent differences between strains in their spore production (Buttery et al. 2009). The strains used in this study likewise showed significant variation in spore production (SMFS: *χ*^2^=3.8e6, df=1, *P*<0.001, GSNP: *χ*^2^=2.9e6, df=1, *P*<0.001, AFCT: *χ*^2^=1.4e6, df=1, *P*<0.001, MLBS: *χ*^2^=4.4e5, df=1 *P*<0.001), allowing us to test this hypothesis for the two sites with significant linearity. To do so, we ranked the strains from most to least dominant based on their clonal spore production (termed ‘prior attribute’ ranking) and compared it to the ranking derived from the outcomes of the pairwise strain interactions (termed ‘observed’ ranking) (Figure 5). If the two rankings are similar, spore production is a robust predictor of social dominance and linearity. Conversely, if the rankings differ substantially, then spore production of the two strains is not a good predictor of social dominance and linearity. Figure 5 shows that in both populations the two rankings exhibited multiple differences in order, suggesting that the differences between strains in their clonal spore production are not strong predictors of the dominance hierarchy.

**Figure 5.**
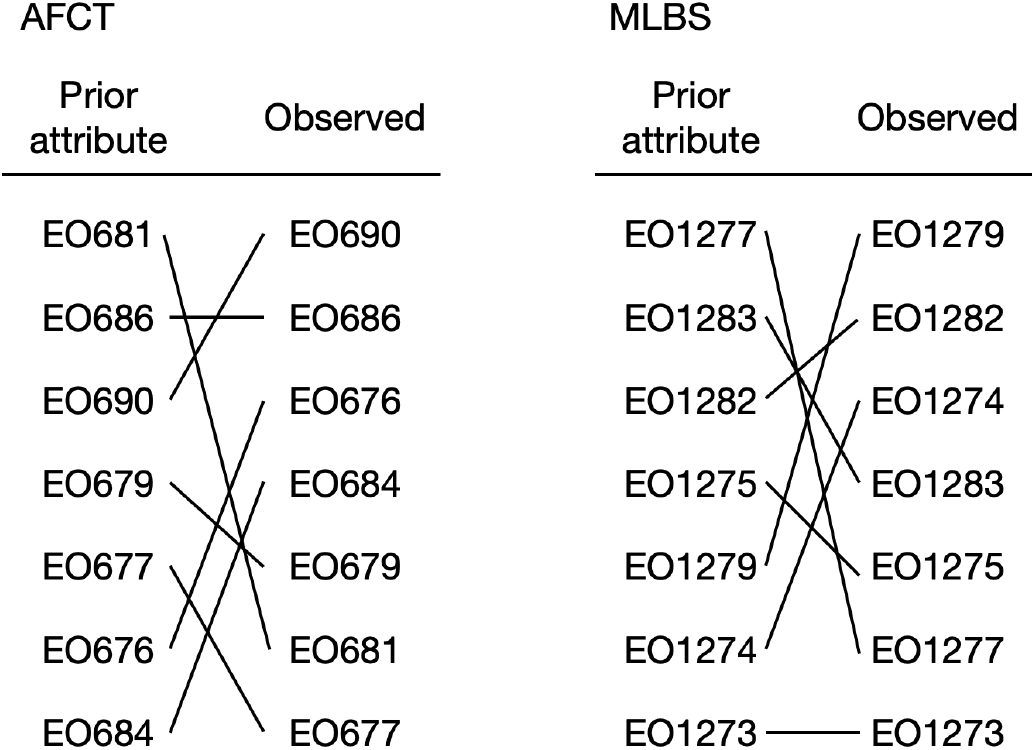
Comparison of the expected dominance rank (left), based on the prior attribute of a strain’s solo spore production, and the observed dominance rank (right) for the two populations (AFCT and MLBS) that exhibit significant linearity. To visualize the changes in the order of the ranking, a single strain in the two rankings is connected by a line, and the crossing of two lines indicates changes in rank order.

### Evidence that prior attributes, direct genetic effects, and social genetic effects influence social dominance

So far, we have examined the variation in social dominance exclusively as a product of a strain’s own predetermined dominance ability—in this case, its ability to sporulate. However, dominance is likely also determined by the traits of the social partner. The relevant traits might not have been identified yet or might only be expressed during the social interaction. The influence of a social partner’s genes on the phenotype of the focal individual is called indirect genetic effects (IGEs) (Wolf et al. 1998). Conversely, direct genetic effects (DGEs) describe the direct influence of an individual’s own genes on its phenotype. In addition, when the interaction between genotypes is dependent on the specific combination of genotypes (i.e., social context) these effects are called genotype-by-genotype (G×G) interactions. Collectively, IGEs and G×G interactions are referred to as the social environment: the environment created by genes present in social partners (Wolf 2003).

To estimate the relative contributions of these different genetic effects to social dominance, we performed a quantitative genetic analysis (Bijma 2014). Since prior work showed that clonal spore production significantly influences a strain’s spore fraction (Buttery et al. 2009), the model also included both the focal and partner strain’s clonal spore production as explanatory factors. Lastly, we included site and genetic distance between strains in a mix. We performed two models: model 1 tested the focal strain’s spore *fraction* in a mix as the response variable, and model 2 tested the focal strain’s *total* spore production in a mix (discussed below).

The results of model 1 showed that a focal strain’s spore fraction in a chimera was significantly influenced by its clonal spore production, as well as that of its partner strain (Table 6). There was no significant effect of genetic distance, and so this term was removed from the final model (χ^2^=0.039, df=1, *P*=0.84). The focal strain’s spore fraction was significantly influenced by the direct genetic effect (DGE) of the focal strain, but not by the indirect genetic effect (IGE) of the partner strain and the interaction between the genotypes of the focal and partner strain (G×G interaction). Analysis of the partitioned genetic effects revealed that 53.1% of the random effects variance was explained by DGEs, 28.5% by IGEs, and 17.4% by G×G interactions (Table 6). Because the focal and partner genotypes can influence dominance through two different terms (their clonal spore production and respective random effects), we assessed their relative contributions using an *R*^2^ metric appropriate for mixed models (Nakagawa and Schielzeth 2013). We found that 11.0% of the total variance could be explained by the fixed effects (i.e., clonal spore production) and 63.7% by the fixed and random effects. Thus, although highly significant, the clonal spore production of the two strains explains only a small fraction of the total variance. This result indicates that the prior attributes were not the main drivers of the outcome: rather, these strains must have other, unidentified attributes that largely determine their ability to dominate the spores.

**Table 6.** The summary of the model that tested the focal strain’s fraction in the spores in chimera as a function of prior attribute clonal spore production and genetic identity of the focal and partner strain and their interaction. The variance explained by the individual genetic effects (DGE, IGE, and G**×**G interaction) and block were calculated by dividing each term’s variance by the sum of the variances of all random effects.

| | Term | df | $\chi^2$ | $P$ | | |
| --- | --- | --- | --- | --- | --- | --- |
| Fixed | Focal clonal spore production | 1 | 11.464 | <0.001 |  |  |
|  | Partner clonal spore production | 1 | 8.551 | 0.003 |  |  |
| | Term | df | $\chi^2$ | $P$ | Variance | % Variance explained |
| Random | Focal genotype (DGE) | 1 | 14.925 | <0.001 | 0.016 | 53.1 |
|  | Partner genotype (IGE) | 1 | 5.268 | 0.022 | 0.009 | 28.5 |
|  | Focal-by-partner interaction (G×G) | 1 | 0.422 | 0.52 | 0.005 | 17.4 |
|  | Block | 1 | 0.036 | 0.85 | 0.0003 | 1.0 |
|  | Total |  |  |  | 0.0303 |  |

We found that total spore production in chimeras is often greater than what is expected based on the average spore production of both strains when developed clonally, consistent with other studies (Buttery et al. 2009; Kuzdzal-Fick et al. 2023). The excess spore production found in this study shows, however, no relationship with the spore inequity observed in that mix (Figure S1; Spearman’s rank correlation: *r=*-0.08, df=76, *P*=0.50). Therefore, modelling variation in a strain’s fraction in the spores as a response variable may ignore important G×G interactions. Specifically, if both strains increase their spore allocation, but do so *equally*, then each strain’s spore fraction would remain at 0.5 despite a strong change in behavior in response to one another. For this reason, we examined not only a strain’s spore *fraction* (i.e., its dominance, as tested in Model 1) in chimera but also its *total* spore production (Model 2).

The results of Model 2 showed that the focal strain’s total spore production in chimera was significantly influenced by the clonal spore production of the partner strain, but not that of the focal strain (Table 7). In addition, there was a significant positive relationship between the genetic distance between the two strains and spore production (estimate=0.003; *P*<0.001; 95% CI: 0.002-0.004). In other words, the more genetically different the two strains are, the more spores the focal strain produces.

**Table 7.**
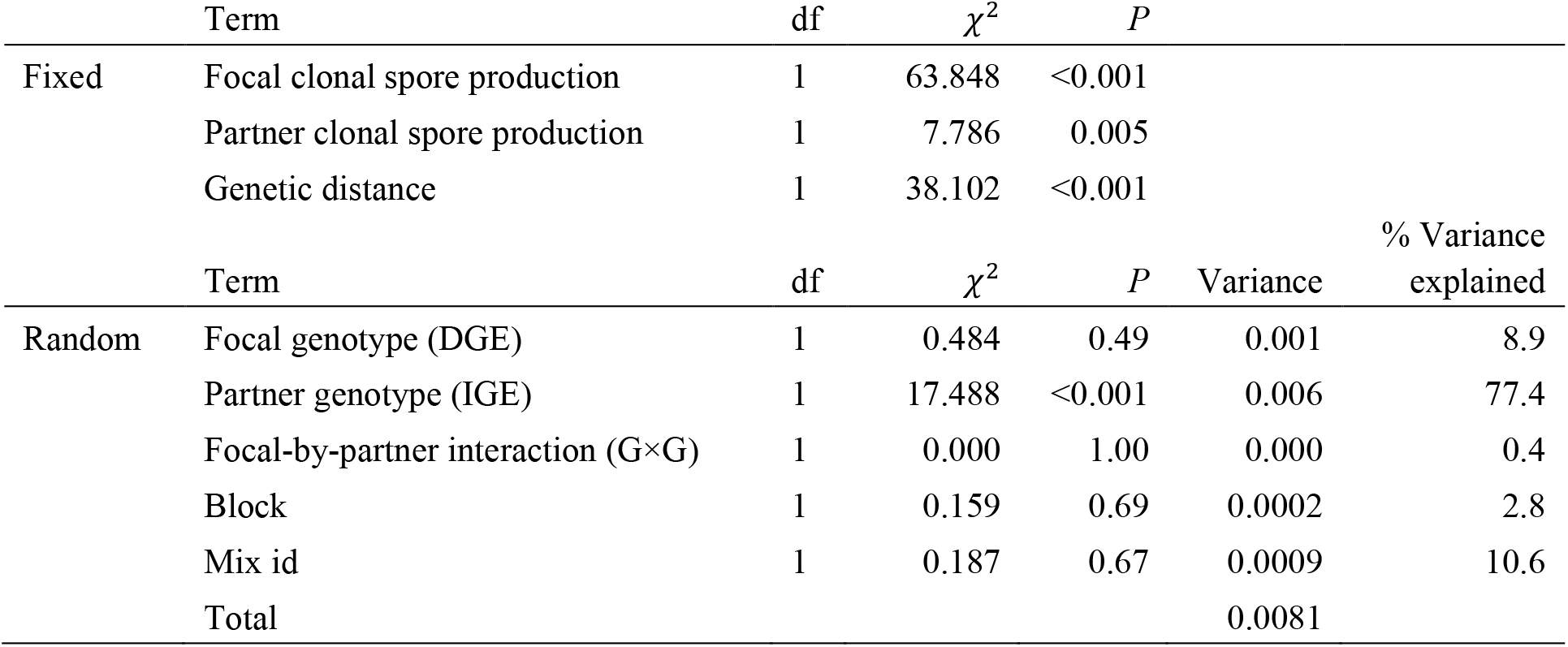
Results of the model that tested the focal strain’s total spore production in chimera as a function of prior attribute clonal spore production, genetic distance, genetic identity of the focal and partner strain, and their interaction. The variance explained by the individual genetic effects (DGEs, IGEs, and G×G interactions) and block were calculated by dividing each term’s variance by the sum of the variances of all random effects.

| | Term | df | $\chi^2$ | *P* | | |
| --- | --- | --- | --- | --- | --- | --- |
| Fixed | Focal clonal spore production | 1 | 63.848 | <0.001 |  |  |
|  | Partner clonal spore production | 1 | 7.786 | 0.005 |  |  |
|  | Genetic distance | 1 | 38.102 | <0.001 |  |  |
| | Term | df | $\chi^2$ | *P* | Variance | % Variance explained |
| Random | Focal genotype (DGE) | 1 | 0.484 | 0.49 | 0.001 | 8.9 |
|  | Partner genotype (IGE) | 1 | 17.488 | <0.001 | 0.006 | 77.4 |
|  | Focal-by-partner interaction (G×G) | 1 | 0.000 | 1.00 | 0.000 | 0.4 |
|  | Block | 1 | 0.159 | 0.69 | 0.0002 | 2.8 |
|  | Mix id | 1 | 0.187 | 0.67 | 0.0009 | 10.6 |
|  | Total |  |  |  | 0.0081 |  |

In contrast to the earlier model, the focal strain’s spore production was significantly influenced by IGEs, but not by DGEs. Similar to before, there was no significant influence of the GxG interactions. Analysis of the partitioned genetic effects revealed that 8.9% of the total variation in spore production was explained by DGEs, 77.4% by IGEs, 0.4% by G×G interactions, 2.8% by block, and 10.6% by mix id (Table 7). We again estimated the *R*^2^ for the fixed and random effects, which showed that 35.7% of the variance in outcome was attributable to the fixed effects, and 44.1% was attributable to the fixed and random effects together. The small difference means that the outcome is largely explained by the fixed effects—the clonal spore production of the two strains and how genetically different they are. While significant, the random effects are small in magnitude. Finally, the total *R*^2^ is also lower compared to the earlier model (44.1% versus 63.7%), indicating that the total spore production of two strains together is less predictable than their percentages.

## Discussion

Here we examined the structure of dominance hierarchies, focusing on their linearity and steepness, in four natural populations of the social amoeba *Dictyostelium discoideum*. In two populations, we identified significantly linear dominance hierarchies. In the third population, the hierarchy was not significantly linear, whereas in the fourth, it was marginally non-linear. Despite this variation, all four hierarchies were shallow, with relatively few significant deviations from spore equity. Together, these results indicate strains with a high degree of competitive similarity for social fitness.

Our finding of less linear and relatively flat social hierarchies (or: *less inequity*) differs from two earlier studies, that used the same seven isolates, which were collected at varying distances over a ∼1 km^2^ area of North Carolina (Fortunato, Queller, and Strassmann 2003; Buttery et al. 2009). Notably, our strains co-occurred in small soil samples of 10 g or less, and the tested strains were no more than 14 cm from each other at the time of sampling—and presumably, often much less. They were thus much more likely to consist of strains that do interact in nature. Our results underscore the importance of an appropriate spatial scale for the study of microbial cooperation in nature.

We expect co-occurring strains to be well-matched for important fitness characteristics, for several reasons. First, a large variance in social fitness can arise when selection is relaxed in the laboratory (Larsen et al. 2023).

By similar logic, low variance is expected for traits subject to strong past selection, and so our observation of little social fitness variation is an important line of evidence that this trait could be experiencing selection in nature. In other words, co-occurring strains might co-occur *because* they do not have large fitness differences. Second, previous studies have shown that spore inequality will drive counter-evolutionary changes to restore fairness (Khare et al. 2009; Hollis 2012; Levin et al. 2015; Miller, Sidell, and Ostrowski 2023). Thus, both ecological and evolutionary processes should promote the co-existence of well-matched competitors.

The level of spore inequity we found in this study is similar in magnitude to what has been observed previously in laboratory screens for cheating behaviors (Santorelli et al. 2008, 2013; Miller, Sidell, and Ostrowski 2023) and another study using a different set of natural strains (Broersma and Ostrowski, in prep.), suggesting that these results are robust and general. In addition, it implies that relatively few genetic differences could underlie this level of spore inequity.

In animal societies, dominance hierarchies are often linear, and much work has been devoted to understanding the determinants of a dominance hierarchy and whether it arises from ‘prior attributes’ (i.e., genetic attributes) or emerges from ‘social dynamics’, which may include winner or loser effects (Chase and Seitz 2011). In this study, we did not allow for winner/loser effects, as we re-grew the strains from frozen for each replicate. That said, the variance attributed to block effects was *more than an order of magnitude smaller* than that attributed to the different genetic effects (Table 6). Overall, these results suggest that the dominance hierarchies, despite being shallow, were repeatable.

In the populations that did exhibit significant linearity, we could investigate the causes of that linearity. We hypothesized that the difference in representation of the strains in the spores could reflect inherent differences in their spore-stalk allocation—their ‘prior attribute’. However, we observed many shifts in rank between the expected performance of strains based on this prior attribute and their actual performance in the mix. Despite this result, the focal and partner strains’ clonal spore production were significant predictors of their percentage of the spores, indicating that these prior attributes do contribute to the establishment of the linear dominance hierarchy, at least in part. Our statistical analyses also indicate that social dominance was significantly influenced by direct and indirect genetic effects, but not genotype x genotype interactions. In addition, we detected these significant genetic effects *after* accounting for the clonal sporulation of the two strains. These results suggest that the linearity results, in part, from additional, non-quantified dominance traits.

A major focus in this paper was on the percentage of the two strains in the spores—the reason for this focus is clear: it reflects *relative* fitness differences between two strains. These differences are paramount for the evolutionary dynamics of allele frequency changes. However, we also considered the *absolute* number of spores produced by each strain. In prior work, we showed that a near-universal response to nonself strains is a facultative increase in the differentiation to spore cells (Kuzdzal-Fick et al. 2023). Moreover, this increase likely results from non-matching *tgrB1-tgrC1* alleles that mediate kin discrimination and influence cell type proportioning responses downstream of this locus (Benabentos et al. 2009; Hirose et al. 2011, 2017; Kuzdzal-Fick et al. 2023). These findings suggested that strains sense nonself and respond selfishly, increasing their investment in the spores relative to the dead stalk. We again observed this response in this study. Taken together, our results are consistent with two types of selfishness, described previously (Liu et al. 2022). On one hand, we see that strains increase their allocation to the spores in the presence of non-identical strains. In the framework of Liu et al., this behaviour corresponds to ‘selfishness’ (i.e., a strategy of cooperating more or less depending on the relatedness of social interactants). On the other hand, we also see significant genetic effects of both the focal and partner strains as determinants of the outcome of a mix. This finding suggests that some of what determines social success may involve manipulative cheating and/or its suppression (Liu et al. 2022).

In summary, our study shows that linearity is indeed a common attribute of social amoeba hierarchies, as determined by four populations at large geographic distances from one another. However, the social amoeba social hierarchies tend to be shallow, indicating low levels of inequity. Nevertheless, there is significant inequity present in natural populations, given that the experimental mixes differed from the clonal controls. We show that prior attributes contribute to the linearity of the dominance hierarchy. However, we also find substantial contributions of DGEs and IGEs, where the traits were not specifically quantified in advance or predicted. Our study adds to a large body of work that indicates social dominance hierarchies are frequently linear. However, the shallowness indicates a relatively fair distribution of the spores among cooperating strains. Whether this substantial cooperation reflects that cheating has not arisen or that it is successfully suppressed is a crucial topic for future study.

## Acknowledgements

The authors would like to thank Smith MacLeish Field Station, Proctor Academy, Mountain Lake Lodge, and Mountain Lake Biological Station for allowing us to collect soil samples. Soil collection in Great Smoky National Park was done under permit GRSM-2017-SCI-2014. We thank Michael Miller, Maria Polo Prieto, Rafael Polo, Jennie Kuzdzal-Fick, Armando Moreno, and Scott Clark for assistance with soil collection and strain isolation. We thank David Fisher for his advice on the statistical models. This work was funded by grants to EAO from the US National Science Foundation DEB-1557023 and the Marsden Fund, administered by Te Apārangi Royal Society of New Zealand, as well as a Massey University Doctoral Scholarship to CMB.

## Author contributions

CMB and EAO conceived the idea and designed the experiments. CMB collected the data. CMB and EAO analyzed the data, wrote the manuscript, and approved the final version.

## Data accessibility

Data and scripts will be made available online after submission.

## Supplementary information

**Table S1.**
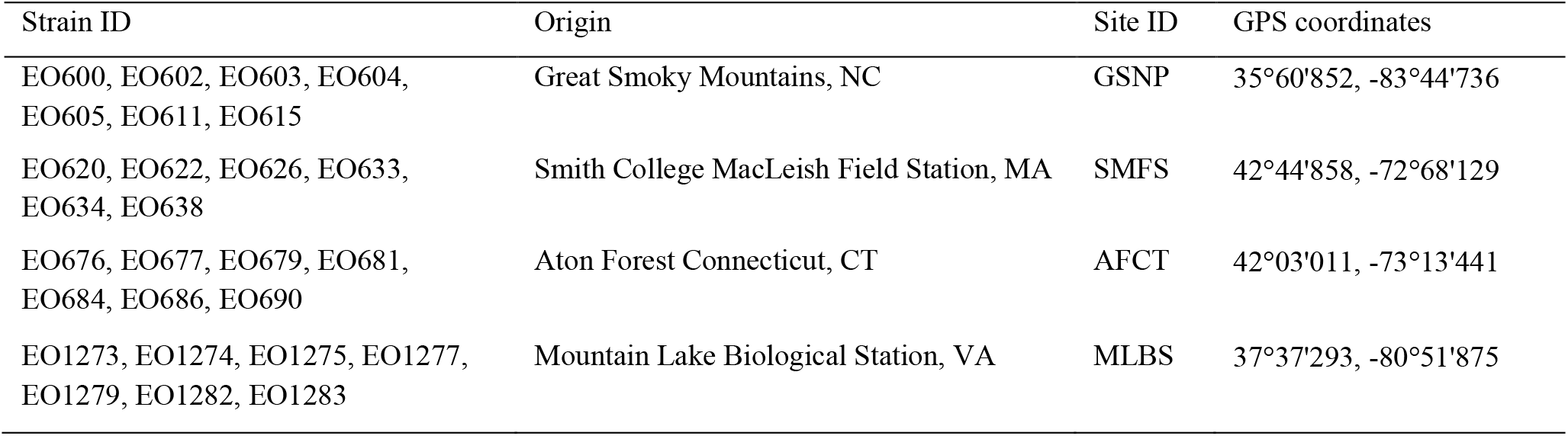
List of strains and their sampling site and GPS coordinates.

**Figure S1.**
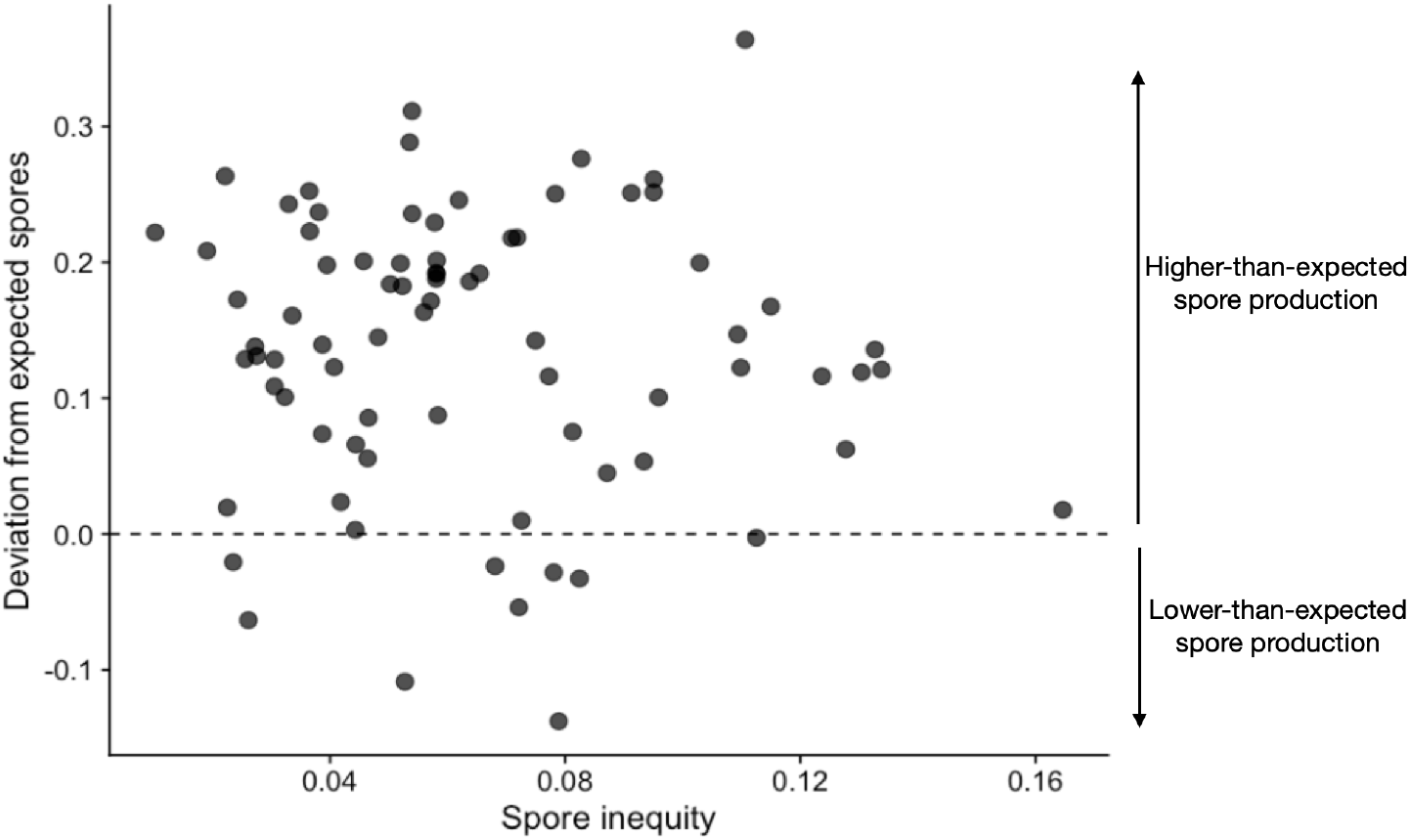
The relationship between the spore inequity and the deviation from the expected total spore production in a mix. Positive and negative values of the deviation from expected spores indicate that a mix produced more or fewer spores than expected based on the average spore production of the strains when developed clonally. 69 out of 78 points lie above the dashed line, indicating that spore production was frequently higher in chimera compared to clonal groups. This increase in total spore production showed however no relationship with the spore inequity observed in a mix, meaning that often both strains increased their spore production to equal numbers.

## Literature cited

Appleby, Michael C. 1983. “The Probability of Linearity in Hierarchies.” Animal Behavior 31: 600–608.

Barkan, C. P. L., J. L. Craig, S. D. Strahl, A. M. Stewart, and J. L. Brown. 1986. “Social Dominance in Communal Mexican Jays Aphelocoma Ultramarina.” Animal Behaviour 34 (February): 175–87.

Barrett, L., S. P. Henzi, T. Weingrill, J. E. Lycett, and R. A. Hill. 1999. “Market Forces Predict Grooming Reciprocity in Female Baboons.” Proceedings of the Royal Society of London. Series B: Biological Sciences 266 (1420): 665–70.

Beaugrand, J. P., and P. A. Cotnoir. 1996. “The Role of Individual Differences in the Formation of Triadic Dominance Orders of Male Green Swordtail Fish (Xiphophorus Helleri).” Behavioural Processes 38 (3): 287–96.

Benabentos, Rocio, Shigenori Hirose, Richard Sucgang, Tomaz Curk, Mariko Katoh, Elizabeth A. Ostrowski, Joan E. Strassmann, et al. 2009. “Polymorphic Members of the Lag Gene Family Mediate Kin Discrimination in Dictyostelium.” Current Biology: CB 19 (7): 567–72.

Bijma, P. 2014. “The Quantitative Genetics of Indirect Genetic Effects: A Selective Review of Modelling Issues.” Heredity 112 (1): 61–69.

Bijma, P., and M. J. Wade. 2008. “The Joint Effects of Kin, Multilevel Selection and Indirect Genetic Effects on Response to Genetic Selection.” Journal of Evolutionary Biology 21 (5): 1175–88.

Buss, L. W. 1982. “Somatic Cell Parasitism and the Evolution of Somatic Tissue Compatibility.” Proc. Natl. Acad. Sci. U S A 79: 5337–41.

Buttery, N. J., D. E. Rozen, J. B. Wolf, and C. R. L. Thompson. 2009. “Quantification of Social Behavior in D. Discoideum Reveals Complex Fixed and Facultative Strategies.” Curr. Biol. 19 (16): 1373–77.

Chase, I. D., and K. Seitz. 2011. “Self-Structuring Properties of Dominance Hierarchies: A New Perspective.” Advances in Genetics 75: 51–81.

Chase, I. D., C. Tovey, D. Spangler-Martin, and M. Manfredonia. 2002. “Individual Differences versus Social Dynamics in the Formation of Animal Dominance Hierarchies.” Proceedings of the National Academy of Sciences of the United States of America 99 (8): 5744–49.

Clutton-Brock, T. H., S. D. Albon, and F. E. Guinness. 1984. “Maternal Dominance, Breeding Success and Birth Sex Ratios in Red Deer.” Nature 308 (5957): 358–60.

Correa, Loreto A., Beatriz Zapata, Horacio Samaniego, and Mauricio Soto-Gamboa. 2013. “Social Structure in a Family Group of Guanaco (Lama Guanicoe, Ungulate): Is Female Hierarchy Based on ‘prior Attributes’ or ‘Social Dynamics’?” Behavioural Processes 98 (September): 92–97.

Curley, J. P. 2016. Compete: Analyzing Social Hierarchies: R Package (version 0.1). https://github.com/jalapic/compete.

Drews, Carlos. 1993. “The Concept and Definition of Dominance in Animal Behaviour.” Behaviour 125 (3): 283–313.

Filosa, M. F. 1962. “Heterocytosis in Cellular Slime Molds.” The American Naturalist 96 (887): 79–91.

Fortunato, A., D. C. Queller, and J. E. Strassmann. 2003. “A Linear Dominance Hierarchy among Clones in Chimeras of the Social Amoeba Dictyostelium Discoideum.” Journal of Evolutionary Biology 16 (3): 438–45.

Goessmann, C., C. Hemelrijk, and R. Huber. 2000. “The Formation and Maintenance of Crayfish Hierarchies: Behavioral and Self-Structuring Properties.” Behavioral Ecology and Sociobiology 48 (6): 418–28.

Hamilton, W. D. 1964a. “The Genetical Evolution of Social Behaviour. I.” Journal of Theoretical Biology 7 (1): 1–16.

Hamilton, W. D. 1964b. “The Genetical Evolution of Social Behaviour. II.” Journal of Theoretical Biology 7 (1): 17–52.

Hausfater, G., J. Altmann, and S. Altmann. 1982. “Long-Term Consistency of Dominance Relations Among Female Baboons (Papio Cynocephalus).” Science 217 (4561): 752–55.

Heinze, J. 1990. “Dominance Behavior among Ant Females.” Die Naturwissenschaften 77 (1): 41–43.

Hirose, Shigenori, Rocio Benabentos, Hsing-I Ho, Adam Kuspa, and Gad Shaulsky. 2011. “Self-Recognition in Social Amoebae Is Mediated by Allelic Pairs of Tiger Genes.” Science 333 (6041): 467–70.

Hirose, Shigenori, Gong Chen, Adam Kuspa, and Gad Shaulsky. 2017. “The Polymorphic Proteins TgrB1 and TgrC1 Function as a Ligand-Receptor Pair in Dictyostelium Allorecognition.” Journal of Cell Science 130 (23): 4002–12.

Hollis, Brian. 2012. “Rapid Antagonistic Coevolution between Strains of the Social Amoeba Dictyostelium Discoideum.” Proceedings. Biological Sciences / The Royal Society 279 (1742): 3565–71.

Huss, Martin J. 1989. “Dispersal of Cellular Slime Molds by Two Soil Invertebrates.” Mycologia 81 (5): 677–82.

Jack, Chandra N., Neil Buttery, Boahemaa Adu-Oppong, Michael Powers, Christopher R. L. Thompson, David C. Queller, and Joan E. Strassmann. 2015. “Migration in the Social Stage of Dictyostelium Discoideum Amoebae Impacts Competition.” PeerJ 3 (October): e1352.

Jackson, Wendy M., and Robin L. Winnegrad. 1988. “Linearity in Dominance Hierarchies: A Second Look at the Individual Attributes Model.” Animal Behaviour 36 (4): 1237–40.

Kessin, R. H. 2001. Dictyostelium. Evolution, Cell Biology, and the Development of Multicellularity. Edited by J. B. L. Bard, P. W. Barlow, and D.L. Kirk. Developmental and Cell Biology. Cambridge University Press.

Khare, Anupama, Lorenzo A. Santorelli, Joan E. Strassmann, David C. Queller, Adam Kuspa, and Gad Shaulsky. 2009. “Cheater-Resistance Is Not Futile.” Nature 461 (7266): 980–82.

Kuzdzal-Fick, J. J., K. R. Foster, D. C. Queller, and J. E. Strassmann. 2007. “Exploiting New Terrain: An Advantage to Sociality in the Slime Mold Dictyostelium Discoideum.” Behavioral Ecology: Official Journal of the International Society for Behavioral Ecology 18 (2): 433–37.

Kuzdzal-Fick, J. J., A. Moreno, C. M. E. Broersma, T. F. Cooper, and E. A. Ostrowski. 2023. “From Individual Behaviors to Collective Outcomes: Fruiting Body Formation in Dictyostelium as a Group-Level Phenotype.” Evolution; International Journal of Organic Evolution 77 (3): 731–45.

Larsen, Tyler J., Israt Jahan, Debra A. Brock, Joan E. Strassmann, and David C. Queller. 2023. “Reduced Social Function in Experimentally Evolved Dictyostelium Discoideum Implies Selection for Social Conflict in Nature.” Proceedings. Biological Sciences / The Royal Society 290 (2013): 20231722.

LeBrun, Edward G. 2005. “Who Is the Top Dog in Ant Communities? Resources, Parasitoids, and Multiple Competitive Hierarchies.” Oecologia 142 (4): 643–52.

Levin, S. R., D. A. Brock, D. C. Queller, and J. E. Strassmann. 2015. “Concurrent Coevolution of Intra-Organismal Cheaters and Resisters.” Journal of Evolutionary Biology 28 (4): 756–65.

Miller, M. E., S. Sidell, and E. A. Ostrowski. 2023. “Costs of Resistance Limit the Effectiveness of Cooperation Enforcement.” BioRxiv. 10.1101/2023.08.29.555259.

Moore, A. J., E. D. Brodie, and J. B. Wolf. 1997. “Interacting Phenotypes and the Evolutionary Process: I. Direct and Indirect Genetic Effects of Social Interactions.” Evolution; International Journal of Organic Evolution 51 (5): 1352–62.

Nakagawa, Shinichi, and Holger Schielzeth. 2013. “A General and Simple Method for ObtainingR2from Generalized Linear Mixed-Effects Models.” Methods in Ecology and Evolution 4 (2): 133–42.

Nakano, Shigeru, and Tetsuo Furukawa-Tanaka. 1994. “Intra and Interspecific Dominance Hierarchies and Variation in Foraging Tactics of Two Species of Stream-Dwelling Chars.” Ecological Research 9 (1): 9–20.

Nelissen, Mark H. J. 1992. “Does Body Size Affect the Ranking of a Cichild Fish in a Dominance Hierarchy?” Journal of Ethology 10 (2): 153–56.

Neumann, Christof, Julie Duboscq, Constance Dubuc, Andri Ginting, Ade Maulana Irwan, Muhammad Agil, Anja Widdig, and Antje Engelhardt. 2011. “Assessing Dominance Hierarchies: Validation and Advantages of Progressive Evaluation with Elo-Rating.” Animal Behaviour 82 (4): 911–21.

Sánchez-Tójar, Alfredo, Julia Schroeder, and Damien Roger Farine. 2018. “A Practical Guide for Inferring Reliable Dominance Hierarchies and Estimating Their Uncertainty.” The Journal of Animal Ecology 87 (3): 594–608.

Santorelli, Lorenzo A., Adam Kuspa, Gad Shaulsky, David C. Queller, and Joan E. Strassmann. 2013. “A New Social Gene in Dictyostelium Discoideum, ChtB.” BMC Evolutionary Biology 13 (January): 4.

Santorelli, Lorenzo A., Christopher R. L. Thompson, Elizabeth Villegas, Jessica Svetz, Christopher Dinh, Anup Parikh, Richard Sucgang, et al. 2008. “Facultative Cheater Mutants Reveal the Genetic Complexity of Cooperation in Social Amoebae.” Nature 451 (7182): 1107–10.

Schjelderup-Ebbe, T. 1922. “Beiträge Zur Sozialpsychologie Des Haushuhns.” Zeitschrift Für Psychologie Und Physiologie Der Sinnesorgane 88: 225–52.

Schjolden, Joachim, Argaudas Stoskhus, and Svante Winberg. 2005. “Does Individual Variation in Stress Responses and Agonistic Behavior Reflect Divergent Stress Coping Strategies in Juvenile Rainbow Trout?” Physiological and Biochemical Zoology: PBZ 78 (5): 715–23.

Schmid, V. S., and H. de Vries. 2013. “Finding a Dominance Order Most Consistent with a Linear Hierarchy: An Improved Algorithm for the I&SI Method.” Animal Behaviour 86 (5): 1097–1105.

Shizuka, Daizaburo, and David B. McDonald. 2012. “A Social Network Perspective on Measurements of Dominance Hierarchies.” Animal Behaviour 83: 925–34.

Smith, J., D. C. Queller, and J. E. Strassmann. 2014. “Fruiting Bodies of the Social Amoeba Dictyostelium Discoideum Increase Spore Transport by Drosophila.” BMC Evolutionary Biology 14 (May): 105.

Strassmann, J. E., Y. Zhu, and D. C. Queller. 2000. “Altruism and Social Cheating in the Social Amoeba Dictyostelium Discoideum.” Nature 408 (6815): 965–67.

Sundström L. Fredrik, Erik Petersson, Johan Höjesjö, Jörgen I. Johnsson, and Torbjörn Järvi. 2004. “Hatchery Selection Promotes Boldness in Newly Hatched Brown Trout (Salmo Trutta): Implications for Dominance.” Behavioral Ecology: Official Journal of the International Society for Behavioral Ecology 15 (2): 192–98.

Valderrabano-Ibarra, C., I. Brumon, and H. Drummond. 2007. “Development of a Linear Dominance Hierarchy in Nestling Birds.” Animal Behaviour 74 (6): 1705–14.

Vehrencamp, S. L. 1983. “Optimal Degree of Skew in Cooperative Societies.” American Zoologist 23 (2): 327–35.

Vries, Han de. 1995. “An Improved Test of Linearity in Dominance Hierarchies Containing Unknown or Tied Relationships.” Animal Behaviour. 10.1016/0003-3472(95)80053-0.

Vries, Han de, Jeroen M. G. Stevens, and Hilde Vervaecke. 2006. “Measuring and Testing the Steepness of Dominance Hierarchies.” Animal Behaviour 71 (3): 585–92.

Vullioud, Colin, Eve Davidian, Bettina Wachter, François Rousset, Alexandre Courtiol, and Oliver P. Höner. 2019. “Social Support Drives Female Dominance in the Spotted Hyaena.” Nature Ecology & Evolution 3 (1): 71–76.

Wilson, E. O., ed. 1975. Sociobiology: The New Synthesis. The Belknap Press of Harvard University Press Cambridge, Massachusetts, and London, England.

Wittemyer, G., and W. M. Getz. 2007. “Hierarchical Dominance Structure and Social Organization in African Elephants, Loxodonta Africana.” Animal Behaviour 73 (4): 671–81.

Wolf, J. B. 2003. “Genetic Architecture and Evolutionary Constraint When the Environment Contains Genes.” Proceedings of the National Academy of Sciences of the United States of America 100 (8): 4655–60.

Wolf, J. B., E. D. Brodie I J. M. Cheverud, A. J. Moore, and M. J. Wade. 1998. “Evolutionary Consequences of Indirect Genetic Effects.” Trends in Ecology & Evolution 13 (2): 64–69.

