## Supplementary information for "Dominance hierarchies are linear but shallow in the social amoeba *Dictyostelium discoideum*"

Table S1. List of strains and their sampling site and GPS coordinates.

| Strain ID | Origin | Site ID | GPS coordinates |
| --- | --- | --- | --- |
| EO600, EO602, EO603, EO604, EO605, EO611, EO615 | Great Smoky Mountains, NC | GSNP | 35°60'852, -83°44'736 |
| EO620, EO622, EO626, EO633, EO634, EO638 | Smith College MacLeish Field Station, MA | SMFS | 42°44'858, -72°68'129 |
| EO676, EO677, EO679, EO681, EO684, EO686, EO690 | Aton Forest Connecticut, CT | AFCT | 42°03'011, -73°13'441 |
| EO1273, EO1274, EO1275, EO1277, EO1279, EO1282, EO1283 | Mountain Lake Biological Station, VA | MLBS | 37°37'293, -80°51'875 |

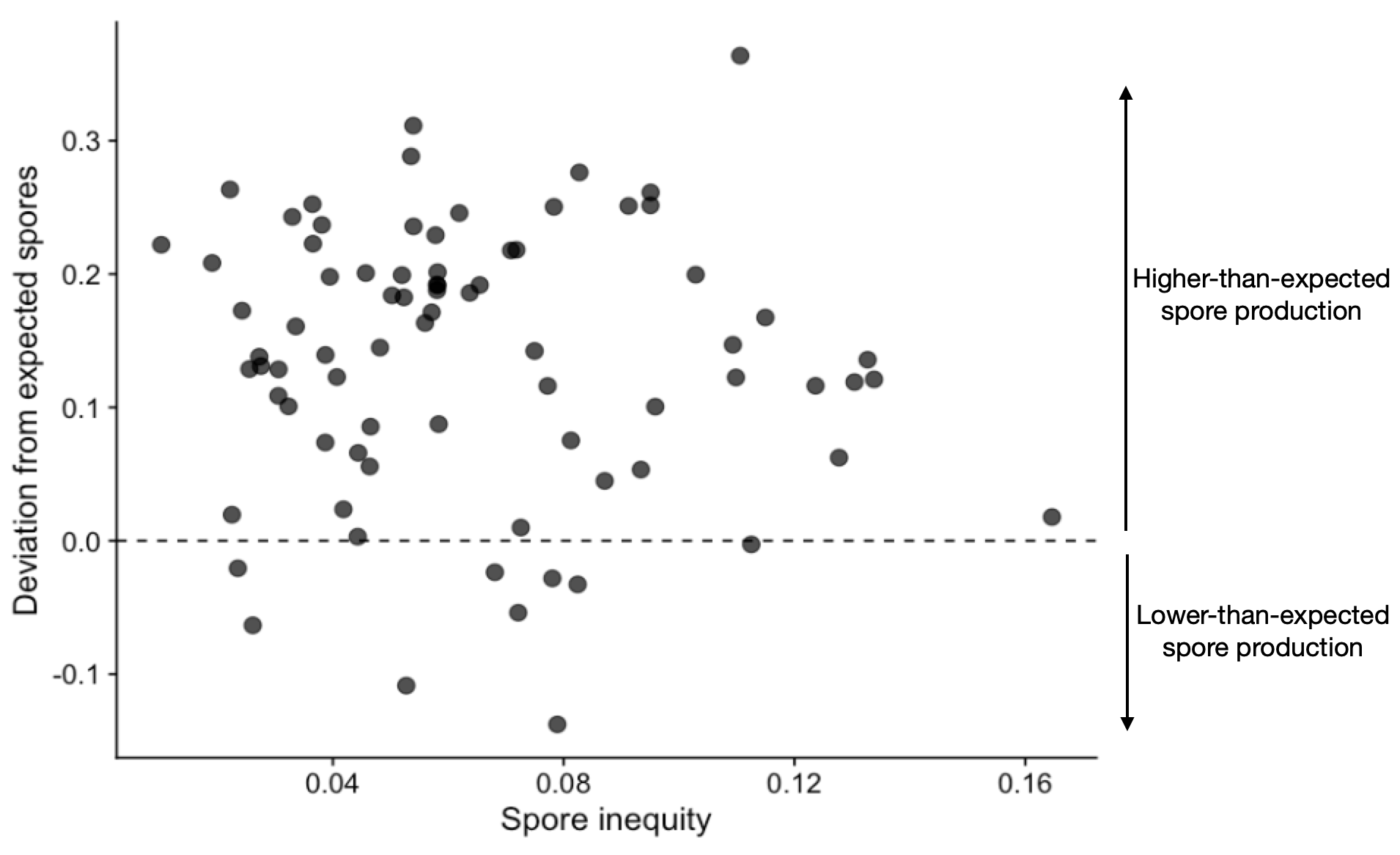

Figure S1. The relationship between the spore inequity and the deviation from the expected total spore production in a mix. Positive and negative values of the deviation from expected spores indicate that a mix produced more or fewer spores than expected based on the average spore production of the strains when developed clonally. 69 out of 78 points lie above the dashed line, indicating that spore production was frequently higher in chimera compared to clonal groups. This increase in total spore production showed however no relationship with the spore inequity observed in a mix, meaning that often both strains increased their spore production to equal numbers.
